# Phase separation versus aggregation behavior for model disordered proteins

**DOI:** 10.1101/2021.06.16.448686

**Authors:** Ushnish Rana, Clifford P. Brangwynne, Athanassios Z. Panagiotopoulos

## Abstract

Liquid-liquid phase separation (LLPS) is widely utilized by the cell to organize and regulate various biochemical processes. Although the LLPS of proteins is known to occur in a sequence dependent manner, it is unclear how sequence properties dictate the nature of the phase transition and thereby influence condensed phase morphology. In this work, we have utilized grand canonical Monte Carlo simulations for a simple coarse-grained model of disordered proteins to systematically investigate how sequence distribution, sticker fraction and chain length influence the phase behavior and regulate the formation of finite-size aggregates preempting macroscopic phase separation for some sequences. We demonstrate that a normalized sequence charge decoration (SCD) parameter establishes a “soft” criterion for predicting the underlying phase transition of a model protein. Additionally, we find that this order parameter is strongly correlated to the critical density for phase separation, highlighting an unambiguous connection between sequence distribution and condensed phase density. Results obtained from an analysis of the order parameter reveals that at sufficiently long chain lengths, the vast majority of sequences are likely to phase separate. Our results predict that classical LLPS should be the dominant phase transition for disordered proteins and suggests a possible reason behind recent findings of widespread phase separation throughout living cells.

## I. INTRODUCTION

Liquid-liquid phase separation (LLPS) of proteins is understood to be a universal biophysical mechanism for the organization and regulation of the intracellular environment.^1–3^ Phase separated assemblies of proteins and RNA/DNA, also known as biological condensates, have been implicated in many key biomolecular processes such as cellular signalling^4^, ribosomal assembly^5^ and transcription of genes^6^. LLPS is often driven by multivalent proteins which act as poylmeric scaffolds that enable the formation of weakly transient networks of noncovalent bonds.^7^ Disorder in protein conformation is also known to play a major role in the formation of these condensates and a large majority of phase separating proteins are known to have intrinsically disordered regions (IDRs).^8,9^ The underlying driving forces include hydrophobic^10^ or electrostatic^11^ interactions and can be regulated by changes in temperature^12^, pH^13^, RNA concentration^14^, salt concentration^15^ as well as the surrounding intracellular environment.

Protein phase separation is highly sensitive to changes in the underlying protein sequence. Performing point mutations at key residue sites is known to disrupt phase separation.^16,17^ Additionally, the sequence patterning of the protein is relevant to its phase separation propensity.^11^ Both analytical theory^18–21^ and explicit chain simulations^22–24^ have been utilized to investigate this sequence-dependent phase behavior. Different sequence-based order parameters such as the sequence charge decoration^25^, sequence hydropathy decoration^26^ and *κ* parameter^27^ have been proposed which correlate the protein sequence and its structural properties (radius of gyration) or phase behavior (critical temperature). In addition to forming through phase separation, biological condensates are also known to form via gelation^28^ or aggregation.^29^ Although the sequence determinants driving protein phase separation have been the subject of extensive investigation, it remains unclear how protein sequence dictates the formation of these alternative phase morphologies, a question of potential significance for both native and de novo engineered condensates.^30^

Recently, highly coarse-grained simulations, in which the protein is modelled as an associative polymer, have emerged as a powerful tool for probing the general principles underlying the sequence dependent phase separation of proteins.^31–35^ Associative polymers can have strongly sequence-dependent phase behavior; depending on their architecture, they may also form a variety of different finite-size aggregates ranging from near-spherical micelles to bilayers, instead of exhibiting classic first order phase separation.^36–39^ Despite the huge diversity of protein sequences, examples of such aggregates maybe quite rare in healthy cells – phase separated condensates appear to be vastly more common.^40^ The reason behind this apparent preponderance of phase separated protein morphologies in biology remains unexplained.

In this work, we have utilized grand canonical Monte Carlo (GCMC) simulations to investigate the connections between protein sequence and the type of phase transition that occurs. GCMC simulations, used alongside standard histogram reweighting techniques, can unambiguously characterize the nature of a phase transition and distinguish macroscopic phase separation from the formation of finite-size aggregates.^41^ Using a coarse-grained lattice model of proteins with purely hydrophobic interactions, we study the influence of sequence composition, patterning and chain length on the nature of the phase transition. By characterizing a data set of 100 model sequences, we show that a suitably normalized sequence charge decoration metric (SCD) works remarkably well at predicting the nature of the transition. For a range of different sequence compositions and chain lengths, we map out the critical value of the normalized SCD and show that phase separation becomes the dominant mode of phase transition for sufficiently long chains. We hypothesize that this size effect could be the reason behind the ubiquity of biological phase separation. Finally, we demonstrate that the normalized SCD is strongly correlated to the critical density, illustrating a fundamental connection between sequence patterning and condensed phase properties.

## II. MODEL AND METHODS

### A. Model for Proteins

In this work, we use a coarse-grained lattice model where the proteins are represented as polymers comprised of two types of entities, hydrophobic/sticky beads which have a net attractive interaction and repulsive hydrophilic beads. Each bead can only occupy a single lattice site and any unoccupied lattice sites are considered to filled by an implicit solvent. Similar lattice models have been extensively used for investigating protein folding and self-assembly.^42,43^ In accordance with conventions used in surfactant literature, we refer to the hydrophobic/sticker segments as “tail” (T) beads and hydrophilic segments as “head” (H) beads. In subsequent figures, hydrophobic beads are represented with red circles and hydrophilic beads with blue circles. Both bonded and non-bonded interactions between neighboring beads can be along the relative position vectors (0,0,1), (0,1,1), (1,1,1) and vectors generated by symmetry operations on this set along the principal axes. This produces a lattice with a coordination number of *Z* = 26. We set the hydrophobic tail beads to have an attractive interaction of *ϵ_TT_* = −1, which also sets the energy scale for the temperature. All other interactions (specifically HH and HT) are set to zero.

### B. Histogram Reweighting Monte Carlo Simulations

GCMC simulations with histogram reweighting were used to investigate the phase behavior of model sequences. Initial runs were performed at a chosen set of temperatures and chemical potentials to obtain the energy and density histograms at those conditions. For a simulation performed at a temperature *T* and chemical potential *μ* in a system with volume *V*, the entropy function at these conditions can be written in terms of the probability of occurrence *f*(*N, U*) of *N* particles with a total energy *U* in the system up to a run specific additive constant *C*.

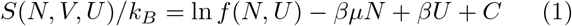

Multiple histograms can be combined using the Ferrenberg-Swendsen^44,45^ method to determine the entropy function of the system across a range of temperatures and chemical potentials. This global entropy function can be utilized to obtain thermodynamic properties of the system at any temperature and chemical potential, given the initial simulation data spanned the range of energies and densities relevant for the new conditions.

### C. Distinguishing phase separation and aggregation

To characterize the nature of the transition, we utilized the system-size dependence of the calculated coexistence curves.^46^ Sequences which undergo a conventional first order phase transition into macroscopic liquid phases have a coexistence curve which is independent of the system size (upper half of Fig. 1). However, for sequences which aggregate, the apparent coexistence curve shows a strong system size dependence (lower half of Fig. 1). Upon increasing the size of the simulation box, there is an apparent decrease in dense phase concentration. This apparent system size dependence can be attributed to the fact that the system forms finite-sized aggregates. Thus, when the system size is increased, the size of the aggregate formed remains unchanged, leading to an apparent reduction in density. This signature of finite aggregation can also be observed from the probability histograms of the density at coexistence as shown in Fig. S1 of the supplementary material.

**FIG. 1.**
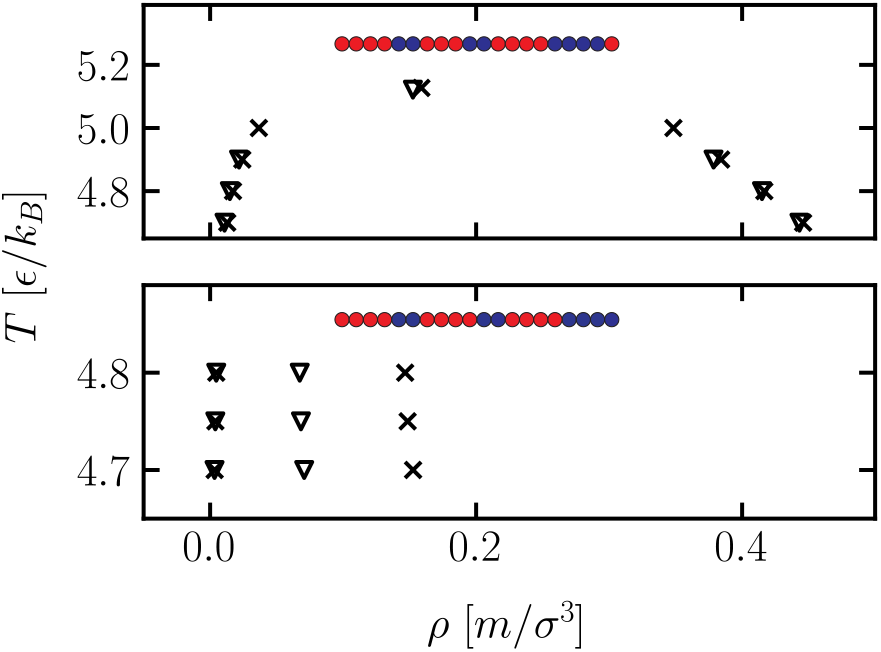
Dense and dilute phase concentrations for *T*_4_*H*_2_*T*_3_*H*_2_*T*_4_*H*_4_*T* (top) and *T*_4_*H*_2_*T*_4_*H*_2_*T*_4_*H*_4_ (bottom) with simulations performed in systems of size *L* = 20*σ* (shown as crosses) and *L* = 30*σ* (shown as inverted triangles). A common density axis is used to highlight the difference in the dense phase concentrations for the two sequences.

We note that aggregation in our model refers to a morphological transformation leading to the formation of a finite aggregate and does not imply irreversibility. Fig. 2 shows snapshots of the dense phase morphology for a phase separating and an aggregating sequence, taken at the same temperature. Importantly, the snapshots illustrate that while the underlying transitions are fundamentally different, there are no major morphological differences between the two dense phases particularly when operating at relatively small system sizes. Thus, it is hard or impossible to distinguish between true phase separation and formation of finite-size aggregates by visual inspection of the simulation box contents alone.

**FIG. 2.**
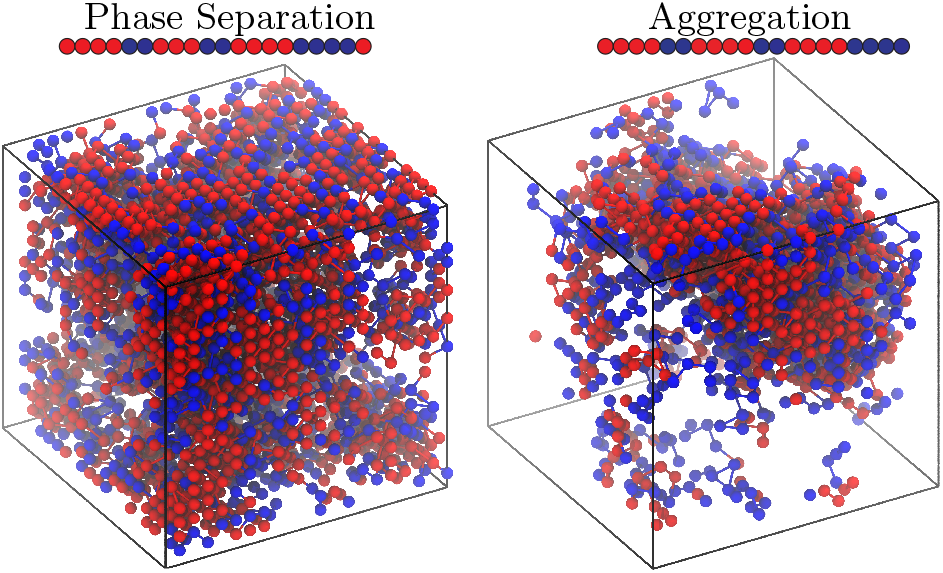
Snapshots of the dense phase morphologies for two sequences, *T*_4_*H*_2_*T*_3_*H*_2_*T*_4_*H*_4_*T* (phase separates) and *T*_4_*H*_2_*T*_4_*H*_2_*T*_4_*H*_4_ (aggregates). Both of these snapshots were taken at a reduced temperature of *T* = 4.8 in a box of size 20*σ* × 20*σ* × 20*σ*. The corresponding phase diagrams for these sequences are shown in Fig. 1. Hydrophobic tail beads are shown in red while hydrophilic head beads are colored blue. The snapshots were generated using VMD.^50^

### D. Estimating critical parameters

For phase separating sequences, we obtained the critical temperature and density using mixed-field finite-size scaling methods.^47–49^ In this approach, an ordering operator 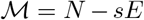 is defined which couples the number of particles *N* to the configurational energy *E* using the field-mixing parameter *s*. For a fixed system size, at criticality, the probability distribution of the scaled ordering parameter 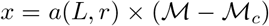 assumes a universal form that depends on the universality class of the underlying first order transition; liquid-liquid phase separation belongs to the three-dimensional Ising universality class. The non-universal parameter *a*(*L, r*) is set to rescale the distribution to unit variance. To obtain the probability distribution of the ordering parameter, we perform a set of GCMC runs near the critical point which are then combined using the Ferrenberg-Swendsen method.^44^ These distributions can be fitted (shown in Fig. 3) to the universal distribution to obtain an estimate for the critical temperature *T_c_* and the critical chemical potential *μ_c_*.

**FIG. 3.**
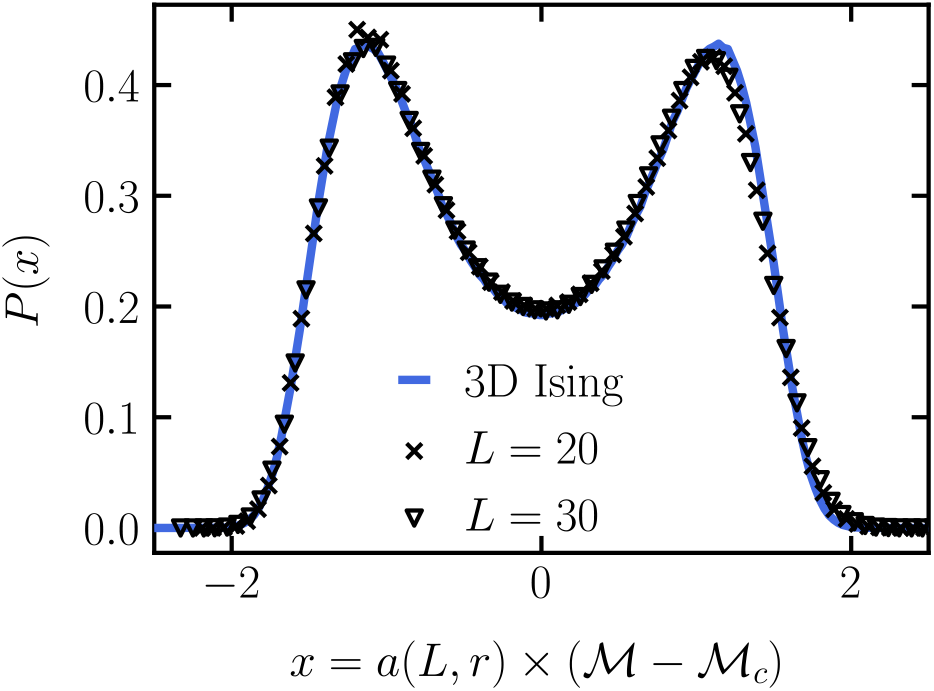
The scaled order parameter distribution from simulation data (shown in symbols) is matched to the universal curve for the 3D Ising universality class (shown in solid line). The sequence used here is *T*_4_*H*_2_*T*_3_*H*_2_*T*_4_*H*_4_*T*, with simulations performed in two different system sizes.

## III. RESULTS AND DISCUSSION

### A. Effects of sequence on phase behavior

To investigate the influence of the polymer sequence on the nature of the transition, we first characterized the phase behavior of chains with a fixed sticker fraction *f_T_* and chain length *r* but having distinct sequence patterning. Five different values of *f_T_* = 0.4, 0.5, 0.6, 0.75, 0.8 were considered with chain length *r* = 20. The sequences studied for this chain length are listed in Table I.

**TABLE I.**
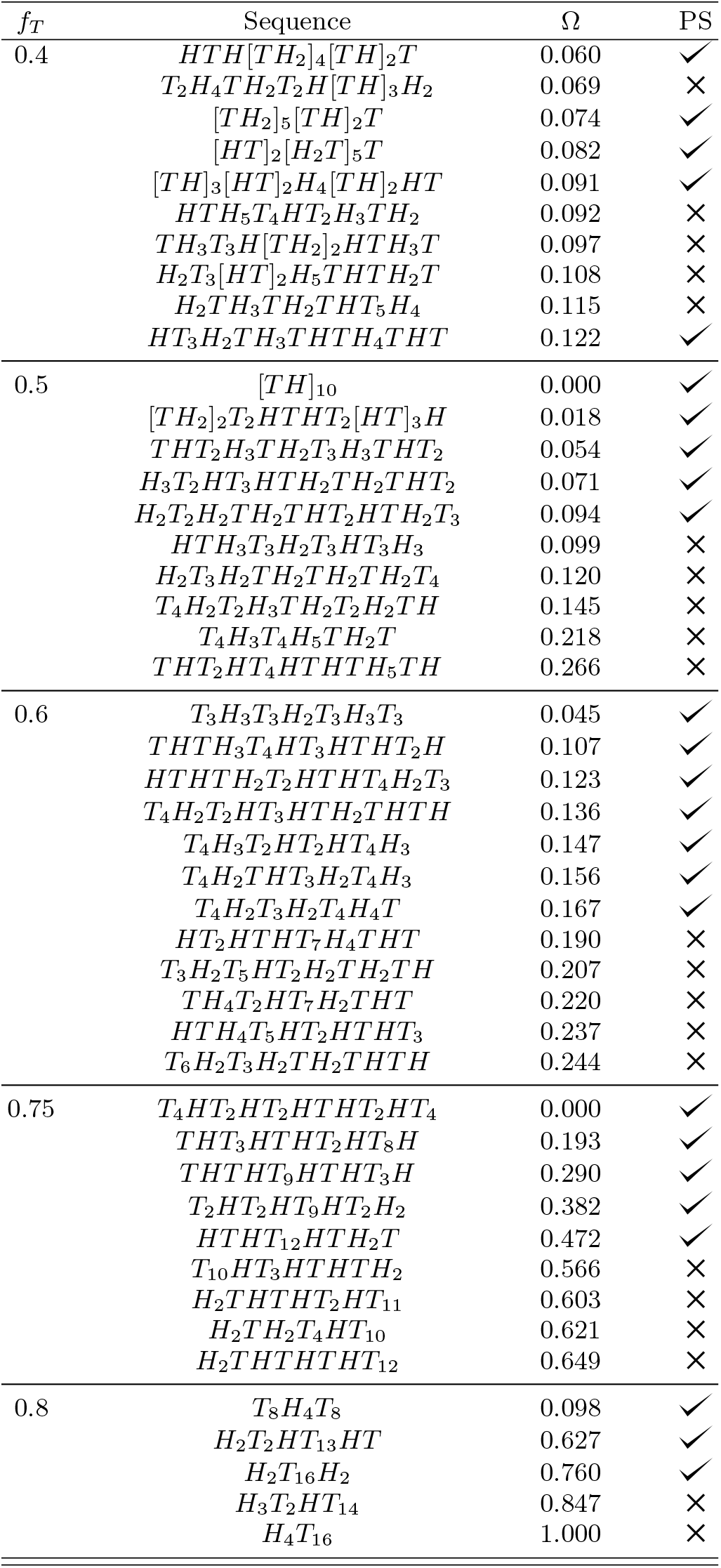
Sequence architecture, sticker fraction, normalized SCD Ω and phase separation capability for sequences of length *r* = 20

We observed that as the degree of dispersion of stickers in the sequence was reduced by clustering them together into longer blocks, the propensity to phase separate decreased. When the dispersion of stickers is reduced beyond a certain point, sequences start showing aggregation behavior and lose the ability to phase separate. These findings are consistent with experimental results which show that phase separation propensity is weakened upon clustering hydrophobic or aromatic “sticky” residues.^51,52^

Furthermore, we found that the transition from phase separation to aggregation depends sensitively on the specific patterning of a sequence. A particularly striking example is seen at *f_T_* = 0.6 for the two sequences *T*_4_*H*_2_*T*_3_*H*_2_*T*_4_*H*_4_*T* and *T*_4_*H*_2_*T*_4_*H*_2_*T*_4_*H*_4_. These two sequences have near identical patterns with the only difference being the position of a single *T* bead. The dilute and dense phase concentrations for these two sequences as a function of temperature are shown in Fig. 1 and snapshots of the dense phase morphologies at *T* = 4.8 are shown in Fig. 2. We also found that among the set of the phase separating sequences for a given (*f_T_, r*), the sequence patterning also influences the critical properties and the shape of the phase envelope (shown in Fig. 4). Additional coexistence data is shown in Fig S2 of the supplementary information.

**FIG. 4.**
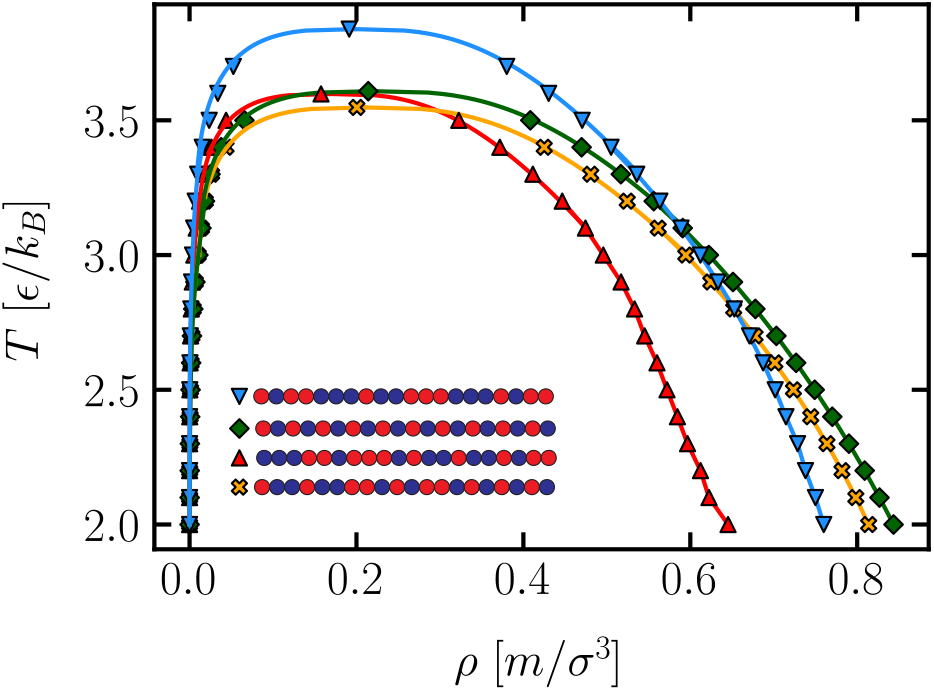
Coexistence curves for sequences with chain length *r* = 20 and sticker fraction *f_T_* = 0.5. The lines are obtained by fitting the near critical coexistence data to the law of rectilinear diameters and the universal scaling relation for densities.^53^

In addition to sequences showing conventional phase separation and aggregation behavior, we also observed that for *f_T_* = 0.6, certain sequences show a “reentrant” transition at low temperatures, with the concentration of the dense phase *decreasing* at lower temperatures, as shown in Fig. 5. Thus, for these reentrant sequences, the condensed phase density reaches a maximum at some intermediate temperature. We find that this anomalous decrease is driven by microphase separation of the hydrophobic sticky blocks at colder temperatures leading to the emergence of voids in the condensed phase; similar behavior has been observed in continuum chain models involving stickers and spacers.^24,34^ In Fig. 6, dense phase morphologies are shown at two different temperatures for the reentrant sequence *T*_4_*H*_3_*T*_2_*HT*_2_*HT*_4_*H*_3_. At *T* = 4.0, the dense phase is observed to be relatively homogeneous with no clear substructure. However, at *T* = 3.0, we see the formation of a lamellar morphology with clear evidence of microphase separation.

**FIG. 5.**
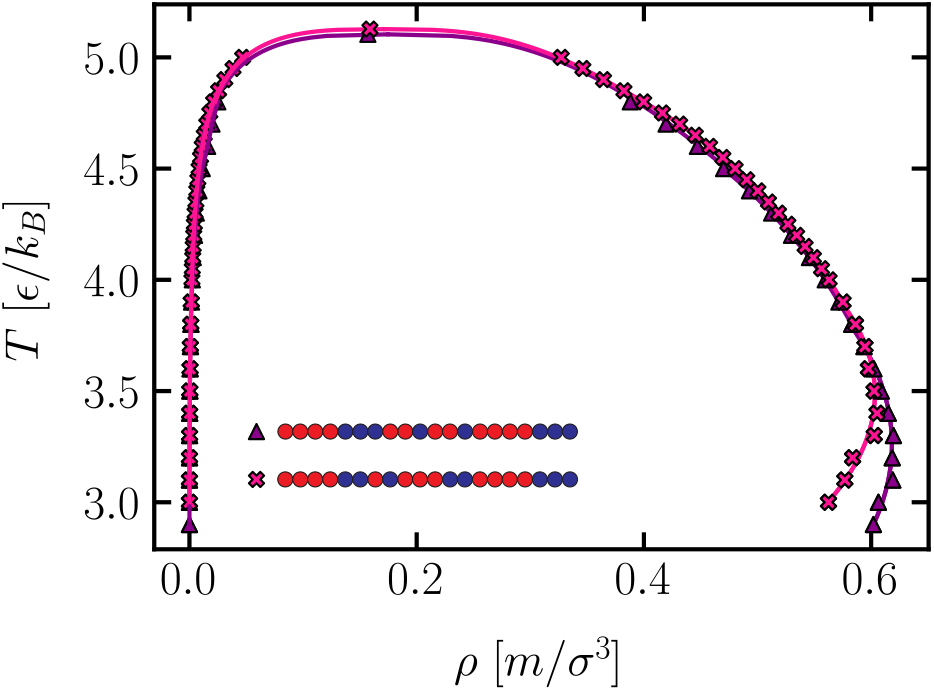
Coexistence curves for reentrant sequences with chain length *r* = 20 and sticker fraction *f_T_* = 0.6. The lines connecting the coexistence points are obtained by fitting the near critical coexistence data to the law of rectilinear diameters and the universal scaling relation for densities. The lines connecting the reentrant points are obtained from a quadratic fit.

**FIG. 6.**
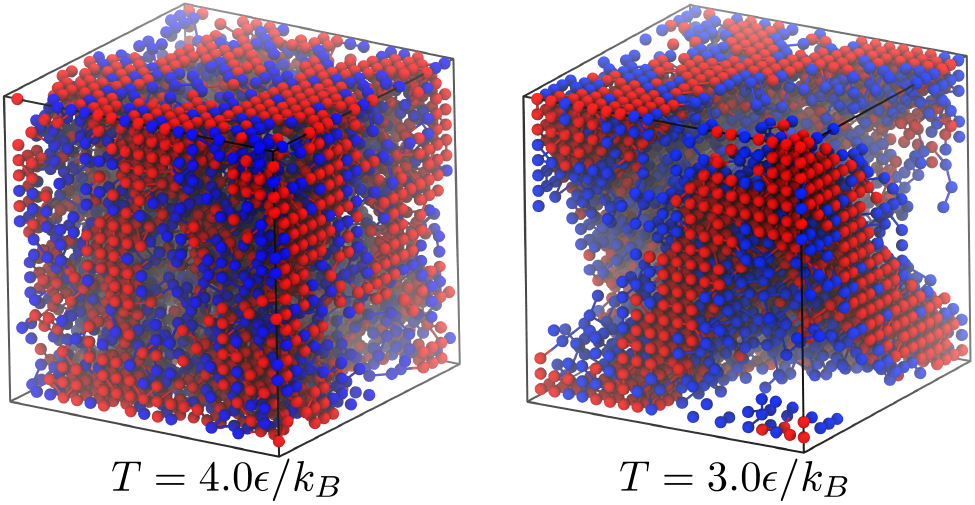
Dense phase morphologies for the reentrant sequence *T*_4_*H*_3_*T*_2_*HT*_2_*HT*_4_*H*_3_ at *T*=4.0*ϵ*/*k_B_* and *T* = 3.0*ϵ*/*k_B_*. These snapshots were taken in a box of size 20*σ* × 20*σ* × 20*σ*. The corresponding phase diagram for these sequences is shown in Fig. 5 (legend symbol: pink cross). The snapshots were generated using VMD.^50^

Recent experimental results have implicated reentrant phase transitions for driving the formation of coreshell type morphologies commonly seen in biological condensates.^54,55^ Although direct analogies cannot be made due to the inherent simplicity of our model, we speculate that the underlying principles might be similar. While it would be interesting to investigate what happens to the density of the condensed phase of these reentrant sequences as we continue to lower the temperature, equilibration becomes extremely difficult at even lower temperatures; thus we restrict ourselves to temperatures at which we are able to equilibrate our systems with certainty.

### B. Normalized SCD: distinguishing phase separation and aggregation

Although there is a clear empirical connection between the sequence patterning and resulting phase behavior, we sought to develop a more quantitative link by establishing a predictive order parameter for the nature of the transition. To do this, we first augmented our data set by further characterizing the phase behavior of sequences with chain length *r* = 40 having sticker fractions *f_T_* = 0.4, 0.5,0.6, 0.8 and *r* = 100 with *f_T_* = 0.5. Data for the sequences studied are shown in Table S1 and S2 in supplementary information. We then tested a set of order parameters previously proposed in protein and polymer literature which have been correlated with structural or condensed phase properties, specifically: 1) sequence charge decoration (SCD)^25^, 2) *κ* parameter^27^, 3) sequence hydropathy decoration^56^ and 4) the mean square fluctuation of block hydrophobicity Ψ^57^.

Among the tested parameters, we observed that the sequence charge decoration (SCD) metric was the one most capable of distinguishing the nature of the transition, performing well across different chain lengths and overall sticker fractions. The SCD was originally developed to capture the effect of charge patterning in polyampholytes and measures the degree of dispersion of a residue in a protein sequence.^25^ In this work we have adapted it to measure the patterning of hydrophobic residues instead. The SCD is defined as:

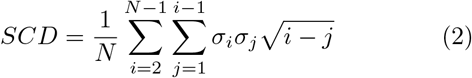

where *N* is the total length of the chain, *i* and *j* refer to positions along the chain and *σ_i_* is the identity of the *i*-th bead. In this work, we have used *σ* = 1 for a hydrophobic bead and *σ* = −1 for a hydrophilic bead. Using this definition, we find that for each (*f_T_, r*) pair, there exists a “soft” threshold value of the SCD beyond which aggregation becomes the dominant behavior.

An undesirable feature of the SCD parameter is that the range of possible SCD values is a strong function of (*f_T_, r*) making global comparisons difficult. To enable comparison across different values of sticker fraction *f_T_* and chain length *r*, we modified the SCD parameter by normalizing it according to the definition:

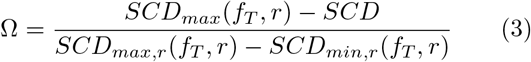

where *SCD_max_*(*f_T_, r*) and *SCD_min_*(*f_T_, r*) are the maximum and minimum possible SCD values for sequences with sticker fraction *f_T_* and chain length *r*. The sequences having the maximum and minimum SCD values are the least and most “blocky” sequences, respectively. This definition simply rescales the value of the SCD between 0 and 1 with the most blocky sequence having Ω = 1 and the least blocky sequence having Ω = 0.

As previously mentioned, even though Ω performs reasonably well as a predictive order parameter, it is not a perfect metric. Ω is invariant with residue inversion i.e changing all *T* beads to H beads and vice-verse will not affect Ω. This becomes prominent for *f_T_* = 0.5 sequences, since every sequence with 50% stickers also has a complement, both of which have identical Ω values. Thus Ω cannot distinguish between a *f_T_* = 0.5 sequence with terminal tail beads and its complement which has terminal head beads. This is problematic because sequences having terminal tail beads are known to have a stronger propensity to phase separate. We observe this near the phase separation threshold where the effect of the terminal beads becomes most pronounced. We also find that Ω has slightly weaker performance at *f_T_* = 0.4 with occasional mispredictions seen even far away from the threshold.

### C. Phase separation thresholds

#### 1. Influence of sticker fraction and chain length

Having established Ω as a soft order parameter, we proceed to define a threshold value for the onset of aggregation. The threshold value Ω* was defined as the average Ω of the two sequences on either side of the transition. Intuitively, we expected that at very low values of the sticker fraction, only the most dispersed chains will be able to phase separate and thus Ω* ≈ 0 for low *f_T_*. Conversely, at high values of *f_T_*, all but the most blocky sequences will phase separate, so Ω* ≈ 1. In Fig. 7 we show that the variation of Ω* with sticker fraction has the expected scaling near the end points. Additionally, for intermediate sticker fractions, we find that the Ω* has a roughly quadratic dependence on *f_T_*.

**FIG. 7.**
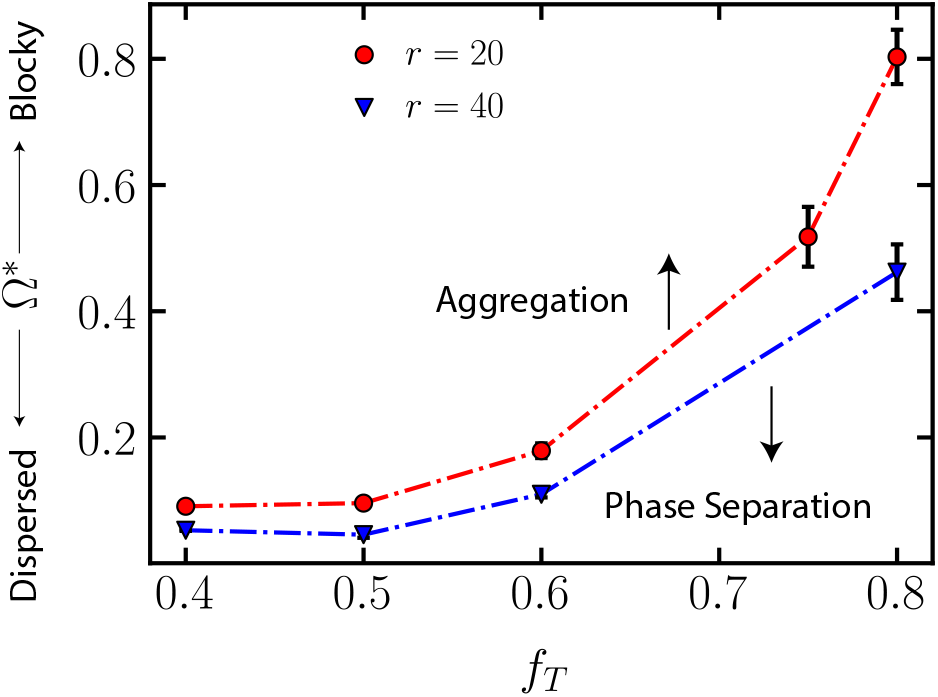
Scaling of the threshold for phase separation Ω* with sticker fraction at a fixed chain length. The dashed lines are meant to guide the eye and demarcate the regimes of phase separation and aggregation for a given chain length. Uncertainties were estimated by measuring differences in the Ω between the two sequences at the boundary between phase separation and aggregation.

The qualitative dependence of Ω* on *f_T_* is robust across chain length for different sticker fractions but the exact dependence of Ω* on chain length remains unclear. To probe this, we computed Ω* for sequences having *f_T_* = 0.5 across chain lengths *r* = 20, 40 and 100. We find that Ω* decreases monotonically with *r* and reaches an asymptotic (non-zero) value as 1/*r* → 0, shown in Fig. 8. From a linear regression, we obtained a Ω* = 0.008 ± 0.003 at the infinite chain limit. Taken together, our results establish a global predictive order parameter for the phase behavior of protein sequences for this model, which is robust across different chain lengths and sticker fractions.

**FIG. 8.**
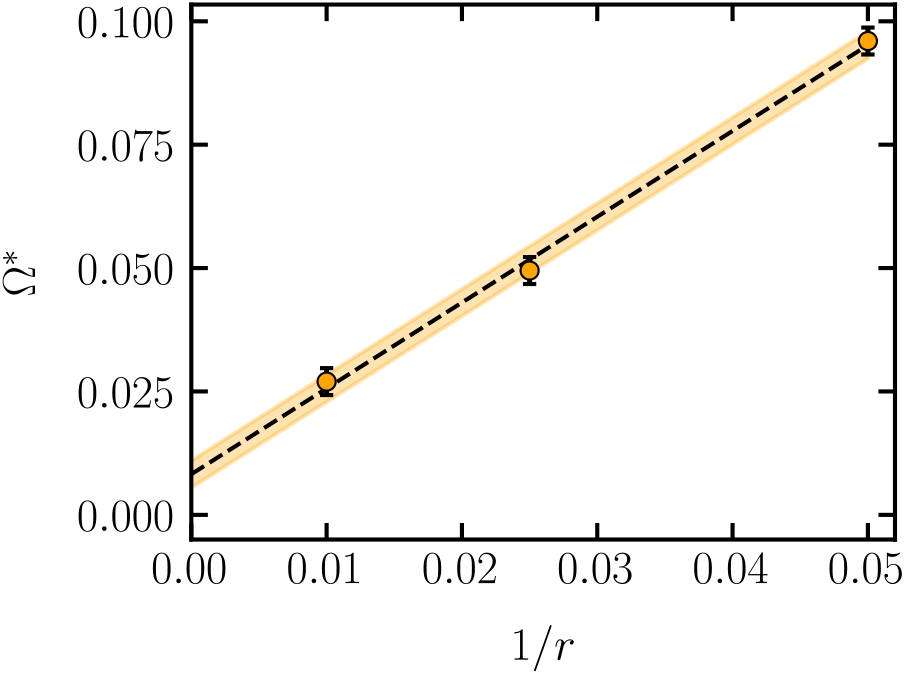
Scaling of the threshold for phase separation Ω* with the inverse chain length for sequences with constant sticker fraction *f_T_* = 0.5. The dotted line represents a linear fit, which is extrapolated to infinite chain length. The shaded area represents the statistical uncertainty measured as the standard error of the fit.

#### 2. Sequence space statistics

Given that we now have an understanding of the threshold Ω = Ω* between phase separation and aggregation, an interesting question to pose is what fraction of possible sequences at a given (*f_T_, r*) lie below the aggregation threshold Ω*. To do this, we first obtained the sequence-space statistics by generating the probability distribution of Ω for different combinations of chain length and sticker fraction. Probability distributions were estimated by generating 10^6^ random sequences for each (*f_T_, r*) and computing the corresponding Ω for each sequence. The fraction of sequences that phase separate was then estimated by integrating the probability distribution up to Ω*.

Fig. 9 shows the probability distributions of Ω for chain length *r* = 40. Vertical arrows in the figure indicate the threshold values Ω* for the different fraction of stickers *f_T_*. The fraction of phase separating sequences increased monotonically with sticker fraction going from 33% at *f_T_* = 0.4 to 78% at *f_T_* = 0.6, in line with the expectation that phase separation should be enhanced with the addition of sticky residues. We then investigated how increasing the length of the chain (at fixed *f_T_* = 0. 5) influences the propensity to phase separate. For *r* = 20, 40 and 100, we compute the fraction of phase separating sequences as 49.9%, 43.6% and 57.0% respectively (Fig. 10). The apparent decrease at *r* = 40 is likely due to inaccuracy in our measurement of Ω*, and we hypothesize that the fraction of phase separating sequences increases monotonically with chain length. To test our hypothesis, we used the chain length dependence of Ω* at *f* = 0.5, as shown in Fig. 8, to obtain the fraction of phase separating sequences at a chain length of *r* = 1000. We find that 97.4% ± 2.3% of all sequences having *r* = 1000 and *f_T_* = 0.5 are expected to phase separate. Thus, there is a very clear increase in the fraction of phase separating sequences for longer chains, with the vast majority of possible sequences capable of phase separating.

**FIG. 9.**
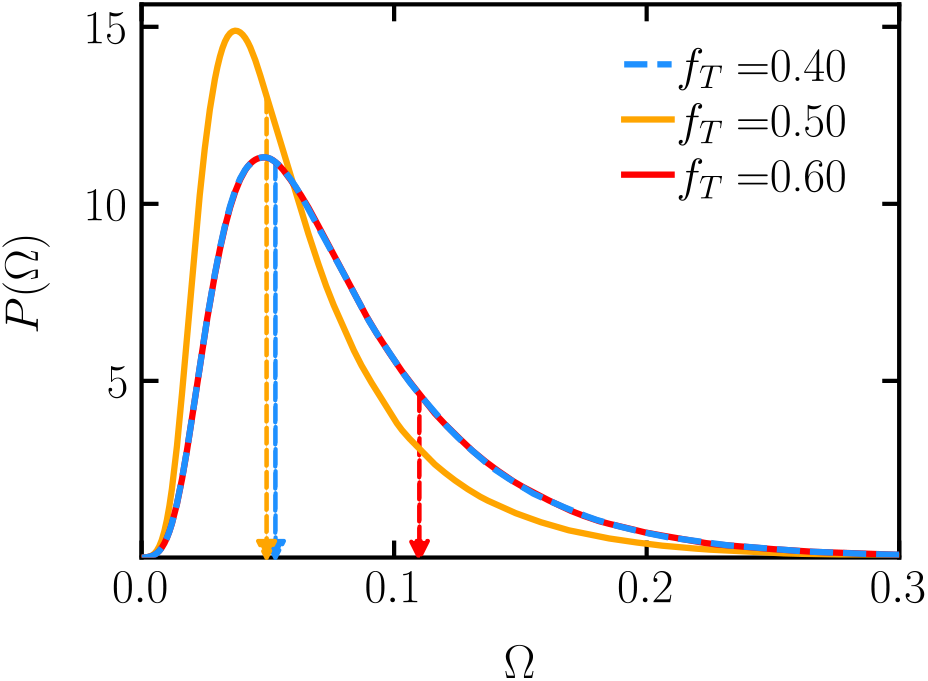
Probability distributions of the normalized SCD Ω for different *f_T_* at chain length *r* = 40. The vertical arrows indicate the threshold value Ω*. Due to the symmetry of Ω, the distributions for *f_T_* = 0.4, shown in blue, and *f_T_* = 0.6, shown in red, are identical but their corresponding Ω* is different.

**FIG. 10.**
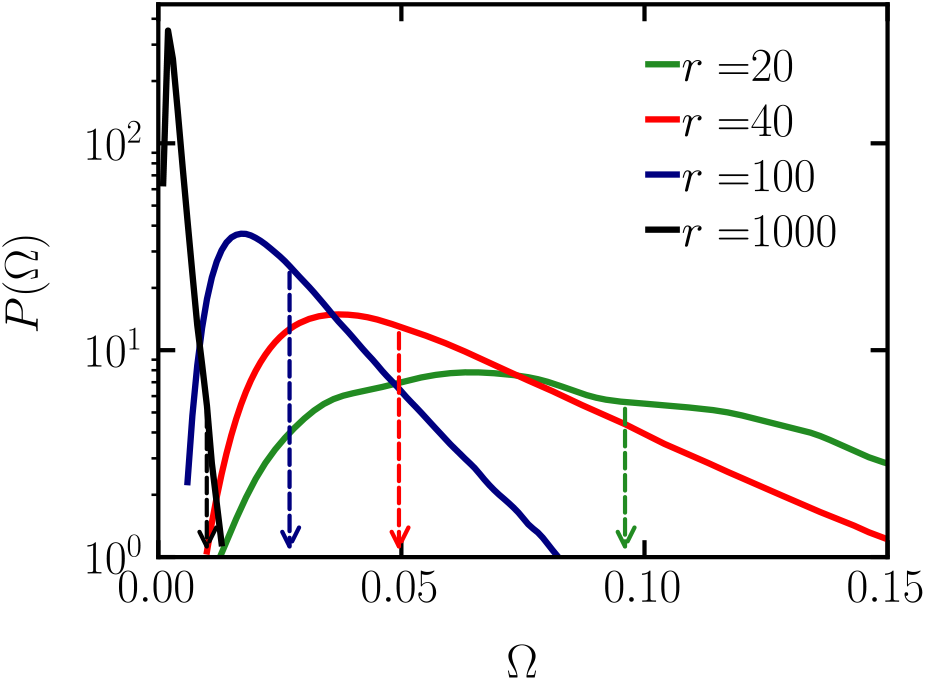
Probability distributions of the normalized SCD Ω for different *r* at sticker fraction *f_T_* = 0.5. The vertical arrows indicate the threshold value Ω*.

This remarkable and unexpected result of a sharp increase in the fraction of phase separating sequences at long chain lengths could have important biological consequences. The ubiquity of biological condensates has been a rather puzzling question. For chains lengths comparable to typical proteins in the cells, our results predict that phase separated morphologies should be overwhelmingly common, with only a tiny fraction of sequences showing aggregation into finite clusters. Our result is consistent with existing experimental evidence and could have important implications regarding the possible phase separation of other long biopolymers such DNA and RNA.

### D. Dependence of critical properties on sequence

In the previous section, we demonstrated that Ω, the normalized SCD parameter, performs well for distinguishing the phase behavior of model sequences. Additionally, for sequences that phase separate, our findings in Sec. III A showed that their coexistence curves were strongly sequence dependent. Here, we investigate whether the dependence of the critical temperature and density on sequence composition and patterning can be rationalized using Ω as the control parameter.

In Fig. 11 and Fig. 12, we show the critical temperatures and densities of sequences having chain length *r* = 20 and 40 with sticker fractions *f_T_* = 0.4,0.5 and 0.6 against the normalized SCD Ω. We find that the critical temperature of sequences is largely decided by the sticker fraction with the precise distribution of stickers in the sequence only seeming to cause small perturbations around this average value. In addition, we also observed sequences at the very edge of phase separation, Ω ≈ Ω*, have a systematically higher *T_c_* than sequences further away from the aggregation threshold.

**FIG. 11.**
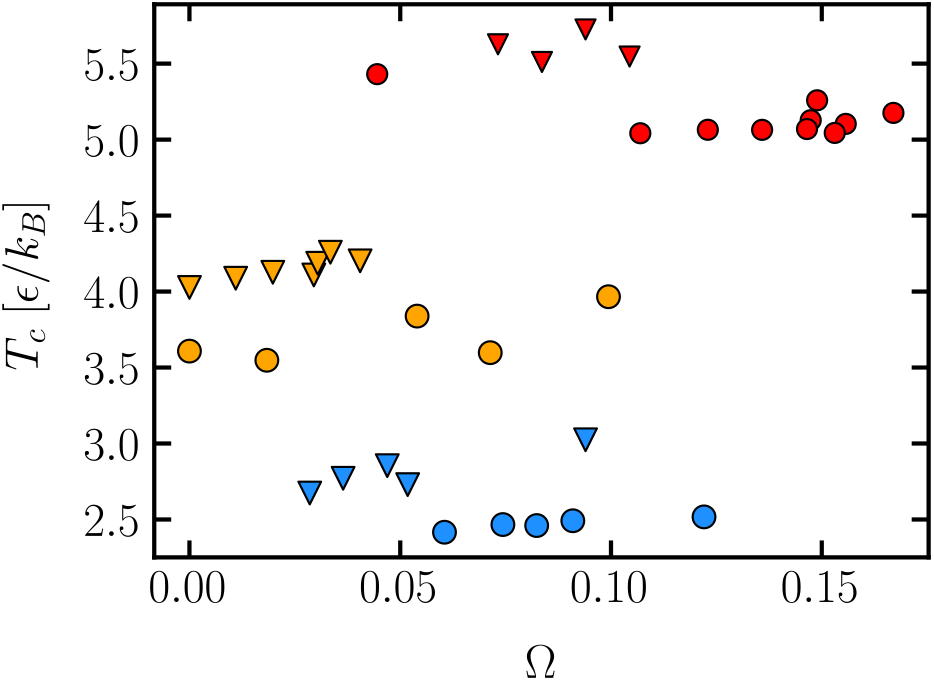
Scaling of the critical temperature with the normalized SCD Ω. The symbol shape represents the chain length: circle *r* = 20, inverted triangle *r* = 40. The color of the symbol is used to represent the fraction of stickers in the chain: *f_T_* = 0. 6 in red, *f_T_* = 0. 5 in orange and *f_T_* = 0. 4 in blue.

**FIG. 12.**
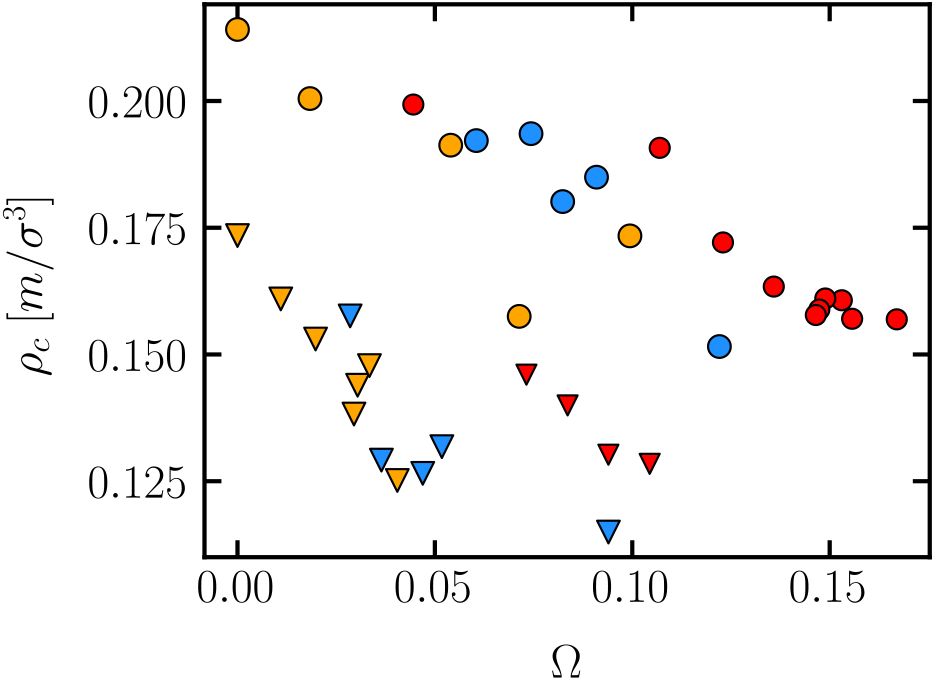
Scaling of the critical density with the normalized SCD Ω. The symbol shape represents the chain length: circle *r* = 20, inverted triangle *r* = 40. The color of the symbol is used to represent the fraction of stickers in the chain: *f_T_* = 0.6 in red, *f_T_* = 0.5 in orange and *f_T_* = 0.4 in blue.

In contrast, the critical density shows a strong negative correlation with the normalized SCD. For both *r* = 20 and *r* = 40, we observe the critical density decreases almost monotonically with Ω until the threshold Ω* is reached. Additionally, for a fixed sticker fraction, the critical density decreases linearly with Ω as seen in Fig. 13. The slope of this line, the fractional change in *ρ_c_* with Ω, increases upon going from *r* = 20 to *r* = 40 and then stays approximately constant upon increasing the chain length further to *r* = 100. Thus we conclude that as the blockiness of the sequence is increased, the density of the condensed phase decreases monotonically until it reaches a minimum value at Ω = Ω*. Below this threshold, the sequence becomes prone to aggregation into finite structures.

**FIG. 13.**
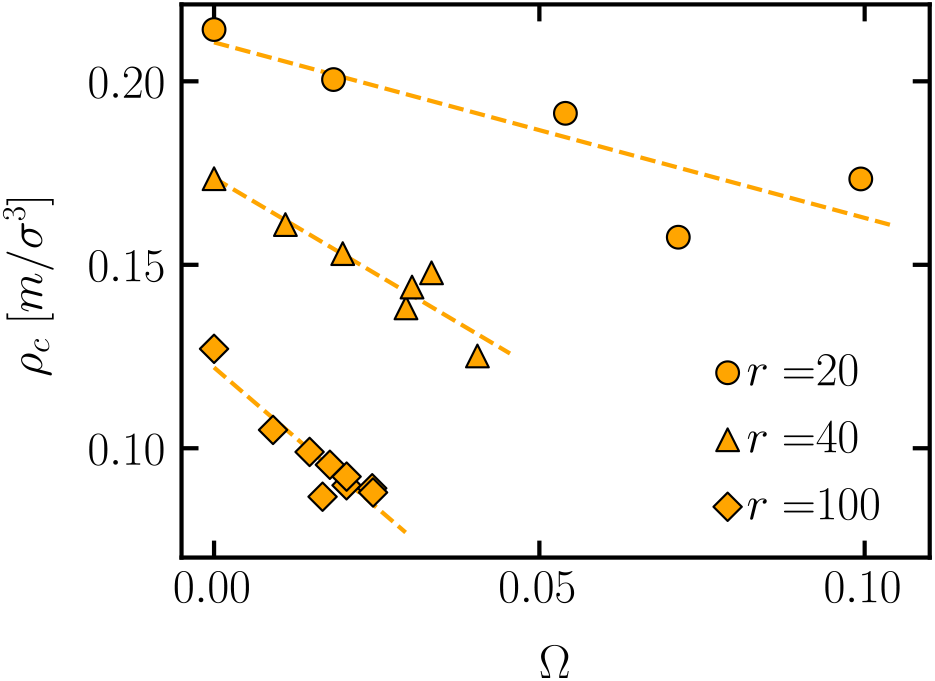
Scaling of the critical density with the normalized SCD Ω for sequences with constant sticker fraction *f_T_* = 0.5 across different chain lengths. The symbol shape represents the chain length.

## IV. CONCLUSIONS

In this work, we have investigated how the sequence patterning of model proteins influences their phase behavior. We found that model proteins can either phase separate or aggregate into clusters of finite extent, depending sensitively on the precise sequence patterning. GCMC simulations combined with histogram reweighting and mapping of a normalized order parameter distribution to the universal Ising curve were found to be sensitive tools to discriminate between phase separation and aggregation and to obtain precise values of the critical parameters. Among the phase separating sequences, we observed that certain sequences exhibit a reentrant transition, with the concentration of protein in the dense phase decreasing as temperature is lowered. This behavior is associated with microphase separation within the condensed phase.

From the characterized phase behavior of 100 different sequences, we found that a normalized sequence charge decoration metric Ω is able to broadly distinguish phase separating from aggregating sequences of the model proteins. Thus, there exists a threshold value Ω*, beyond which the ability to phase separate into a macroscopic phase is lost and sequences become aggregation prone. Although we have focused on the relation between sequence blockiness and finite-size aggregation in this work, experiments also suggest a potential link between clustering of residues and the propensity of forming irreversible protein aggregates.^58^ Further theoretical and experimental efforts will be needed to investigate this connection.

Using the normalized SCD Ω, we found that at a constant chain length, the threshold normalized SCD Ω* has an approximately quadratic dependence on the sticker fraction (hydrophobicity) *f_T_*. At a constant sticker fraction, our results show that Ω* scales linearly with inverse chain length and reaches an asymptotic non-zero value at infinite chain length. Since the Ω is intrinsically related to the overall blockiness of the sequence, our result establishes a robust connection between blockiness in the sequence patterning and its underlying phase behavior. In addition to hydrophobic patterning, charge patterning is also known to play an important role in driving protein LLPS. However, unlike hydrophobic residues, clustering of charges is seen to enhance phase separation tendency.^11^ Investigating the cumulative effects of charge and hydrophobic patterning will be necessary to develop a complete picture of sequence dependent protein phase behavior.

To estimate what fraction of sequences of a certain length and sticker fraction are likely to phase separate, we obtained the sequence space statistics by calculating the distribution of Ω for a given (*r*, *f_T_*) and utilized this distribution. Our results show that the fraction of phase separating sequences increases monotonically with sticker fraction at a constant chain length. The variation with chain length was found to be nearly monotonic with a relatively minor change in the fraction of phase separating sequences when going from chain length *r* = 20 to *r* = 100. However, for *r* = 1000, we found a dramatic increase in the fraction phase separated, with 98% of possible sequences predicted to phase separate.

From our results, we conclude that the phase separation propensity increases rapidly as a function of chain length. Our findings suggest that at sufficiently long chain lengths, the vast majority of possible sequences will phase separate irrespective of sticker fraction or sequence patterning. We hypothesize that the ubiquity of biological phase separation may simply be tied to the fact that most biologically relevant proteins are sufficiently long to be in the regime where phase separation becomes dominant. This would also explain why finite aggregation behavior is relatively rare in biology despite the huge diversity of possible protein sequences.

## ACKNOWLEDGMENTS

This research was primarily supported by the Princeton Center for Complex Materials (PCCM), a U.S. National Science Foundation Materials Research Science and Engineering Center (Grants No. DMR-1420541 and DMR-2011750). The authors thank Antonia Statt for valuable comments and discussions. Simulations were performed using computational resources provided by the Princeton Institute for Computational Science and Engineering (PICSciE) and the Office of Information Technology’s High Performance Computing Center and Visualization Laboratory at Princeton University.

## DATA AVAILABILITY STATEMENT

The data that support the findings of this study are available from the corresponding author upon reasonable request.

## V. SUPPLEMENTARY MATERIAL

### A. Distinguishing phase separation and finite aggregation from density histograms

**FIG. S1.**
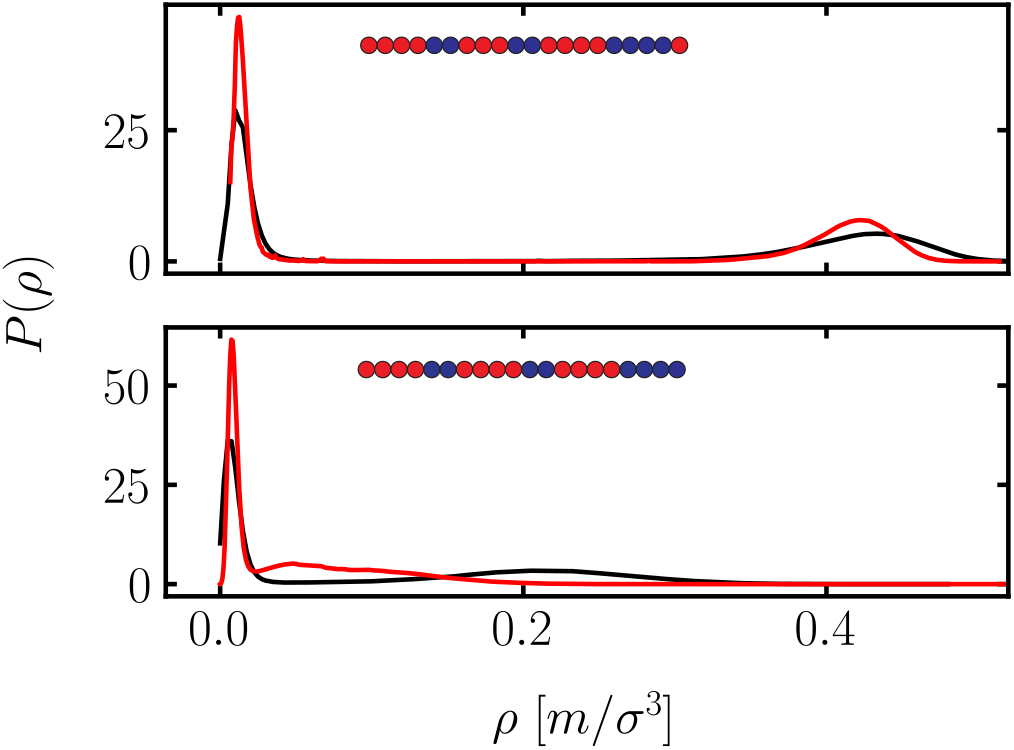
Histograms of the density at *T* = 4.8 and coexistence chemical potential for *T*_4_*H*_2_*T*_3_*H*_2_*T*_4_*H*_4_*T* (top) and *T*_4_*H*_2_*T*_4_*H*_2_*T*_4_*H*_4_ (bottom) with simulations performed in systems of size *L* = 20*σ* (black) and and *L* = 30*σ* (red). For the phase separating sequence *T*_4_*H*_2_*T*_3_*H*_2_*T*_4_*H*_4_*T*, the dilute and dense peaks are invariant with system size. In contrast, there is a shift in the location of the dense phase peak for aggregating sequence *T*_4_*H*_2_*T*_4_*H*_2_*T*_4_*H*_4_.

### B. Phase diagrams of sequences with *f_T_* = 0.6

**FIG. S2.**
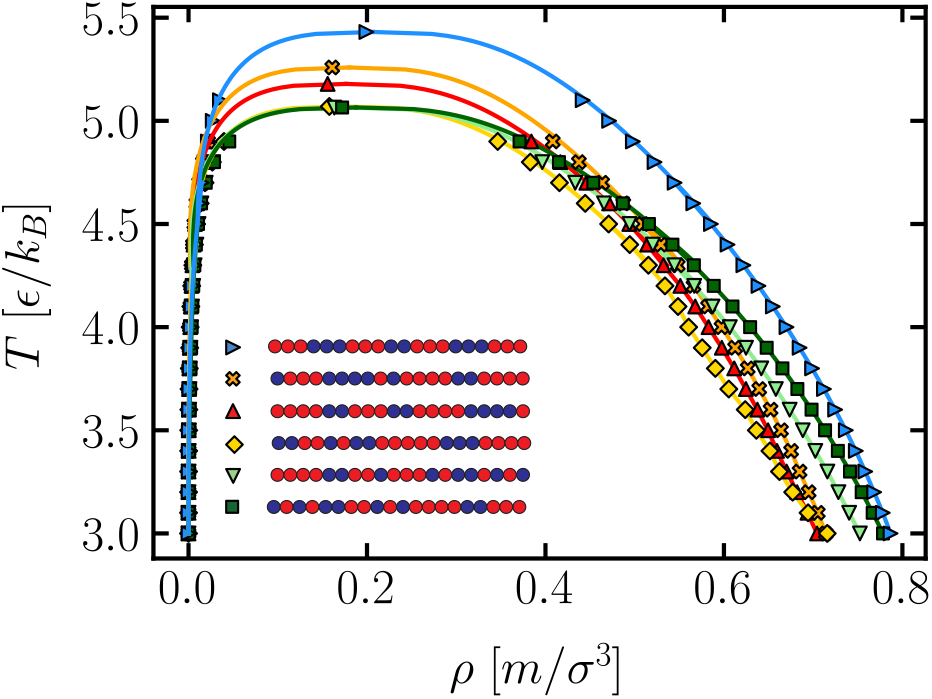
Coexistence curves for sequences with chain length *r* = 20 and sticker fraction *f_T_* = 0.6. The lines are obtained by fitting the near critical coexistence data to the law of rectilinear diameters and the universal scaling relation for densities.

**TABLE S1.**
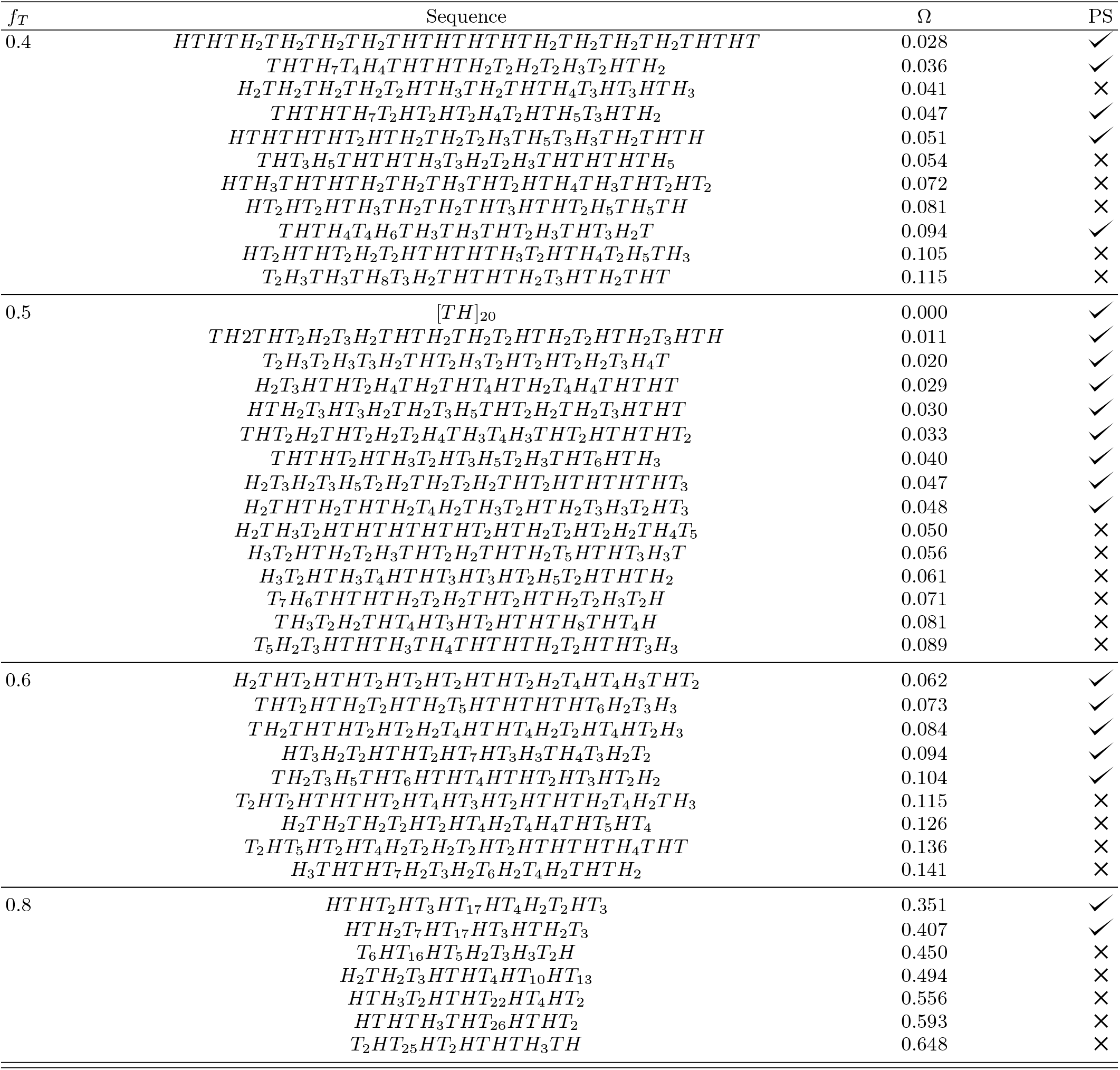
Sequence architecture, sticker fraction, normalized SCD Ω and phase separation capability for sequences of length *r* = 40

**TABLE S2.**
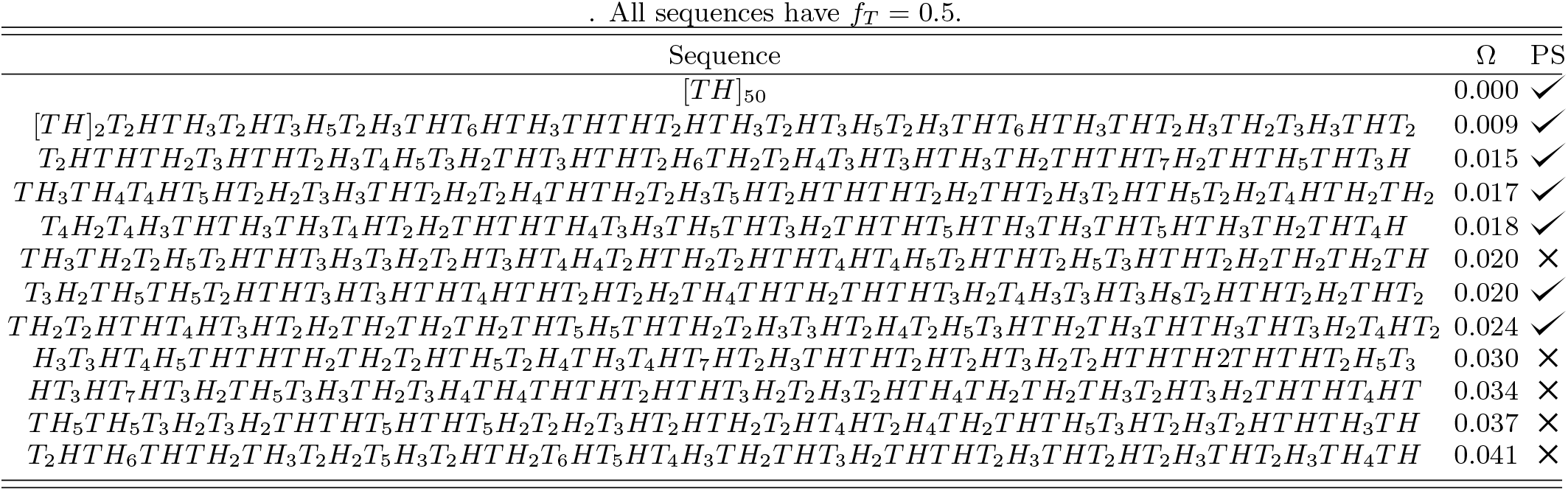
Sequence architecture, sticker fraction, normalized SCD Ω and phase separation capability for sequences of length *r* = 100

